# Association of the gut microbiota with colorectal cancer in a South Asian cohort of patients

**DOI:** 10.1101/694125

**Authors:** AGP Sahankumari, Bawantha D. Gamage, Gathsaurie Neelika Malavige

**Author notes:** Corresponding author: Dr. Bawantha Gamage, MBBS,MRCS,MS,FAMASI,FCSSL, Department of Surgery, Faculty of Medical Sciences, University of Sri Jayewardanepura, Nugegoda, Sri Lanka, P.O. Box 10250.

## Abstract

**Background:** As the gut microbiome is thought to play a role in the pathogenesis of colorectal carcinoma (CRC) and affected by the diet and the genetic composition, we sought to investigate the patterns of gut microbiota that associate with CRC in a South Asian cohort of patients with CRC.

**Methodology:** The relative abundance of 45 types of gut microbial species were determined in faecal samples of CRC patients (n=24), DM (n=20) and healthy age matched controls (n=44), using a PCR array. Data was analyzed using the specific software for analysis of bacterial DNA quantification.

**Results:** The species *Bacteroides fragilis* (23.9-fold), *Bacteroides thetaiotaomicron* (8-fold) and *Akkermansia muciniphila* (5.9 fold) were several-fold over expressed in patients with CRC compared to healthy individuals, whereas bacterial species of the Phylum Proteobactria were under expressed. There was no difference in the abundance of these 3 species of bacteria with tumour stage or gender and age of patients. Aeromonas species, *Enterococcus faecium* and *Shigella dysenteriae* (Proteobacteria) were over 100-fold over abundant in those with DM compared to healthy individuals. Although 70.83% of those with CRC also had diabetes, the relative abundance of microbiota in CRC patients were different to those who had diabetes and no CRC.

**Conclusions:** Patients with CRC and DM harbor a markedly different gut microbiota patterns compared to their healthy counterparts. Similar patterns of gut microbial dysbiosis that associate with CRC and DM appear be seen in South Asian populations, compared to Western countries, despite differences in the diet and ethnicity.

## Background

Colorectal cancer (CRC) is the third commonest cause of cancer worldwide and is the fourth commonest cancer leading to death [1]. It has been predicted that the deaths due to colonic cancer and rectal cancer will increase by 60% and 71.5%, respectively until year 2035 due to the increase in the aging population [1]. The increase in the incidence of CRC is predicted to rise substantially more in developing countries vs developed countries due to these changes in population demographics[2]. As this increase in the incidence of CRC in developing countries is likely to result in a huge burden to their economies, there is an urgent need to implement programs that reduce its occurrence and adopt novel diagnostic and treatment methods of CRC.

Diet and lifestyle are major risk factors for development of CRC along with genetic susceptibility, the presence of metabolic diseases such as diabetes and obesity and inflammatory bowel disease [3-6]. Metabolic diseases and CRC are associated with microbial dysbiosis, which is characterized reduced diversity of the gut microbiome with an overabundance of the genera Proteobacteria and Firmicutes [7]. Several types of microbiota have been shown to associate with CRC such as pks-positive E.coli, enterotoxigenic *Bacteriodes fragilis, Fusobacterium nucleatum* and *Streptococcus gallolyticus* [8-12]. While some of these microbes were overabundant in the gut microbiome of patients with CRC, some have been detected specifically in tumor tissue and also in distance metastasis, suggesting that they may play a role in the pathogenesis of this cancer [12]. They are thought predispose to the development of CRC by inducing epigenetic changes and thereby affecting gene transcription, inducing DNA damage and reactive oxygen species and by inducing procarcinogenic cytokines [12].

Of the factors that affect the diversity of the gut microbiome, the diet plays a central role. Although the relative abundance of gut microbiota depends on an individual’s genetic composition (12%), the influence of the diet is much greater (57%)[13]. Individuals who predominantly consume a Mediterranean diet, rich in grains, legumes, nuts, vegetable and fruits were found to have a gut microbiome which reduced the risk of metabolic diseases, inflammatory bowel disease and colonic cancer compared to those who consume a typical Western diet [14]. South Asian individuals have a very different diet than those of Western and the South East Asian populations, due to differences in religious and cultural practices. Their diets are typically rich in grains, pulses, vegetables and fruits with a low intake of red meat. These vast differences in the diet are likely to influence the microbial composition and thus protect or predispose to the development of CRC. In fact, is has been shown that the gut microbiome is significantly different in individuals of different ethnicity, living in the same geographical area [15]. Since dietary factors are likely to directly contribute to the microbial composition and thus to the risk of developing CRC, change in the dietary patterns can be an important strategy in the prevention and treatment of CRC[16]. Therefore, in order to implement such preventive and therapeutic strategies, it would be important to initially characterize the gut microbial patterns in South Asian individuals with CRC living in those countries.

In this study, we have determined the relative abundance of 45 species of gut microbiota on patients with CRC, aged matched healthy individuals and also in patients with metabolic diseases such as diabetes. We found that that pattern of the gut microbiota was significantly different in those with CRC and diabetes compared to healthy individuals.

## Methods

### Patients

We recruited 24 patients with CRC who underwent colonoscopy at Colombo South Teaching Hospital, Sri Lanka between January 2017 and April 2018, following informed written consent. Stool samples were obtained from these patients two weeks after colonoscopy. All clinical details regarding altered bowel habits, abdominal pain, loss of weight, appetite along with laboratory and radiological investigations such full blood count, ultrasound scanning of the abdomen and CT scans were recorded and CRC grading was carried out according to TNM staging classification [17].

### Recruitment of healthy controls and patients with diabetes mellitus

In order to compare the changes that associate with CRC, we recruited healthy individuals (n=44) who underwent colonoscopy and were found not to have any bowel pathology and who were non-obese, (BMI <23.9), waist circumference <80cm for females and <90cm for males and who did not have diabetes or hyperlipidemia. Again, the stool samples were obtained from these patients two weeks after colonoscopy.

In addition to the above controls, as most of the patients with CRC also had diabetes mellitus (DM), in order to differentiate the changes in the patterns of gut microbiota with those that are specific to CRC, we also recruited patients with DM (n= 20) who underwent colonoscopy and were found not to have any CRC or gut pathology, between January 2017 to April 2018. As in patients with CRC, stool samples were obtained from these individuals, two weeks following colonoscopy. Patients who had adenomas, who received antibiotic therapy for more than 1 week prior to stool sample collection, who received chemotherapy and/or radiation or with a history of CRC and inflammatory bowel disease were excluded.

### Ethics Statement

All subjects provided informed written consent prior to participating in the study. Ethical approval was granted by Ethical Review Committee, Faculty of Medical Sciences, University of Sri Jayewardenepura (Application No: 35/16).

### Sample collection and DNA Extraction from stools

The stool samples were collected two weeks after the patients underwent colonoscopy giving time for the gut microbiota to re-establish itself. The stool samples were transported to the laboratory within 24 hours after collection. DNA was extracted using QIAamp DNA Stool Mini Kit (QS, Hilden, Germany) according to the manufacturer’s instructions and the extracted DNA stored at - 20°C prior to quantification of the stool microbiota.

### Quantification of the gut microbiota

The Microbial DNA qPCR Array Intestinal Infection 2 kit (Qiagen, Hilden, Germany) (Supplementary table 1) was used to amplify species specific 16S rRNA genes in order to identify and to quantify the relative abundance of 45 types of gut microbiota according to the manufacturer’s instructions (Qiagen, Hilden Germany) (Supplementary table 1). Briefly, 5000ng (0.005 mg) extracted bacterial DNA from each stool specimen was used in the microbial qPCR master mix and target-specific fluorescent probes. The reaction was performed in an *Applied Biosystems7500*, 96-well plate detection system. qPCR cycling condition first step comprise of one cycle of initial PCR activation step for 10 minutes at 95°C, following 40 cycles of denaturation for 15 sec at 95°C and annealing and extension for 2 minutes at 60°C as the second cycling reaction step. The threshold cycle value (Ct) for each reaction was determined by manually setting the threshold limit. The relative abundance of different bacterial species was determined by comparing the Ct value of each bacterial species with the Ct values of same bacterial species generated from the healthy controls using the Baid 1407 intestinal infections microbial profiling data analysis software (Qiagen, Hilden Germany).

### Statistical analysis

Statistical analysis was performed using Graph PRISM version 7. Differences in the relative abundance of different bacterial species in patients with CRC, patients with diabetes and healthy individuals were compared using the Mann-Whitney U test (two tailed),

## Results

### Characteristics of patients with CRC

Of the 24 patients with CRC, 15 (62.5%) were males and 9 (37.5%) were females. The median age of those with CRC was 59 years (IQR 53.25 to 64 years), in those with DM was 65 (IQR 59.5 to 68 years) and in healthy individuals was 53 (IQR 47 to 65 years). In patients with CRC, 4/24 (16.67%) of the tumors were present in the descending colon, 4/24 (16.67%) in the sigmoid colon, 11/24 (45.83%) in the rectum, 2/24 (8.33%) in recto –sigmoid junction and 1/24 (4.17%) in hepatic flexure, 1/24 (4.17%) in anal verge and 1/24 (4.17%) in transverse colon. 17/24 (70.83%) of those with CRC had DM.

2/24 of CRC patients were in stage 0 and i based on the TNM classification at the time of diagnosis, 7 were in stage ii, 8 s in stage iii and 7 patients were in stage iv. None of the patients with CRC, DM or the healthy individuals had a previous history of inflammatory bowel disease or CRC. The clinical details of all patients with CRC is shown in Table 1.

**Table 1:**
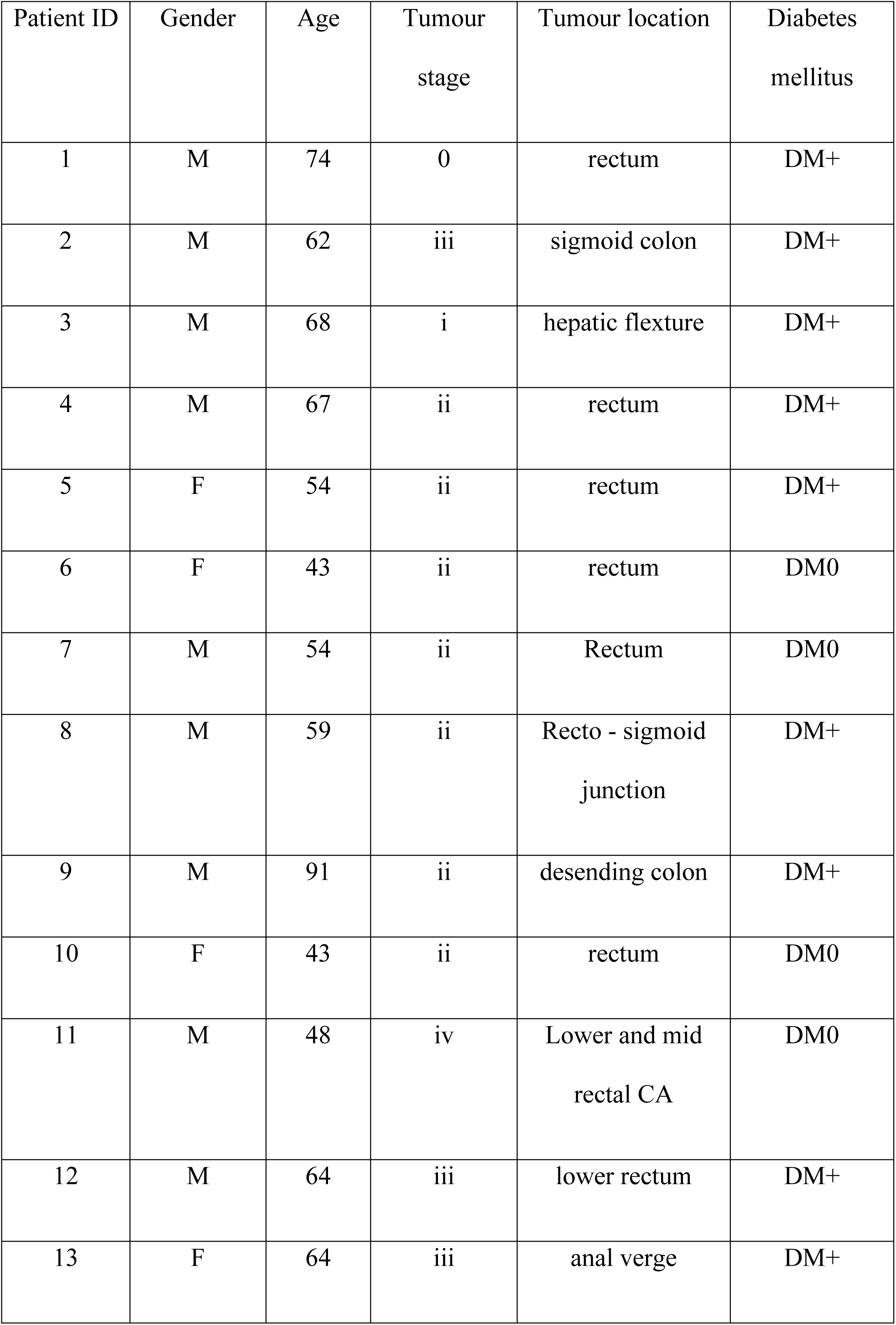
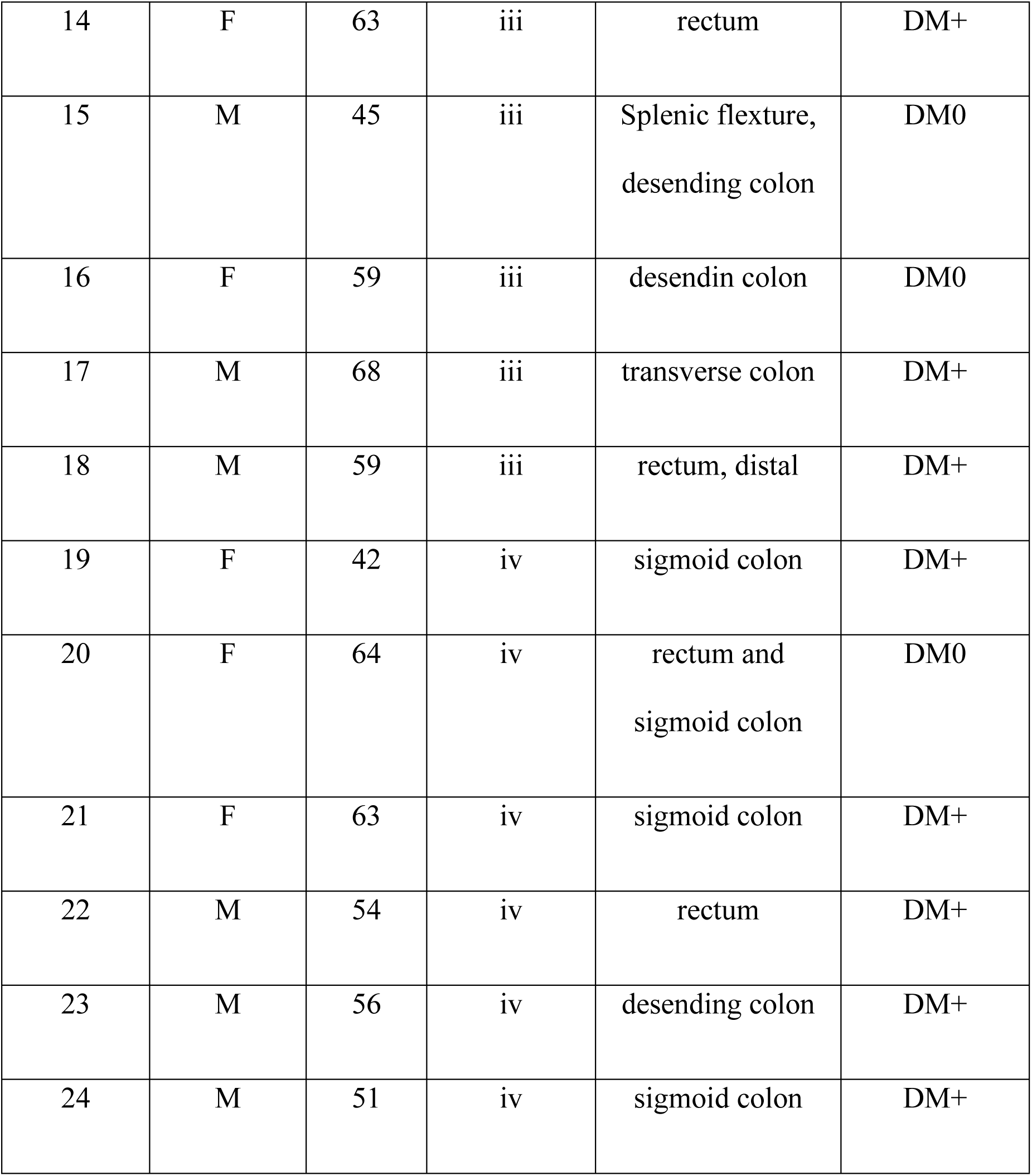
Clinical characteristics of patients with Colorectal cancer.

### Patterns of gut microbiota in CRC compared to healthy individuals

As the relative abundance of gut microbiota has shown to be markedly different in those with CRC compared to healthy individuals^7^, we proceeded to determine the patterns of gut microbiota in our cohort of patients, in comparison to healthy individuals.

We observed marked differences in the gut microbiota patterns in patients with CRC when compared to healthy individual groups (Fig 1). The five most abundant bacterial species among in patients with CRC compared to healthy individuals were *Bacteroides fragilis, Bacteroides thetaiotaomicron, Akkermansia muciniphila, Aeromonas spp. (Aeromonas enteropelogenes, Aeromonas hydrophila, Aeromonas punctata, Aeromonas media*) and *Bacteroides vulgatus* (Table 1). Most notably *Bacteroides fragilis* was 23.9-fold over expressed in those with CRC compared to healthy individuals followed by *Bacteroides thetaiotaomicron*.

**Figure 1:**
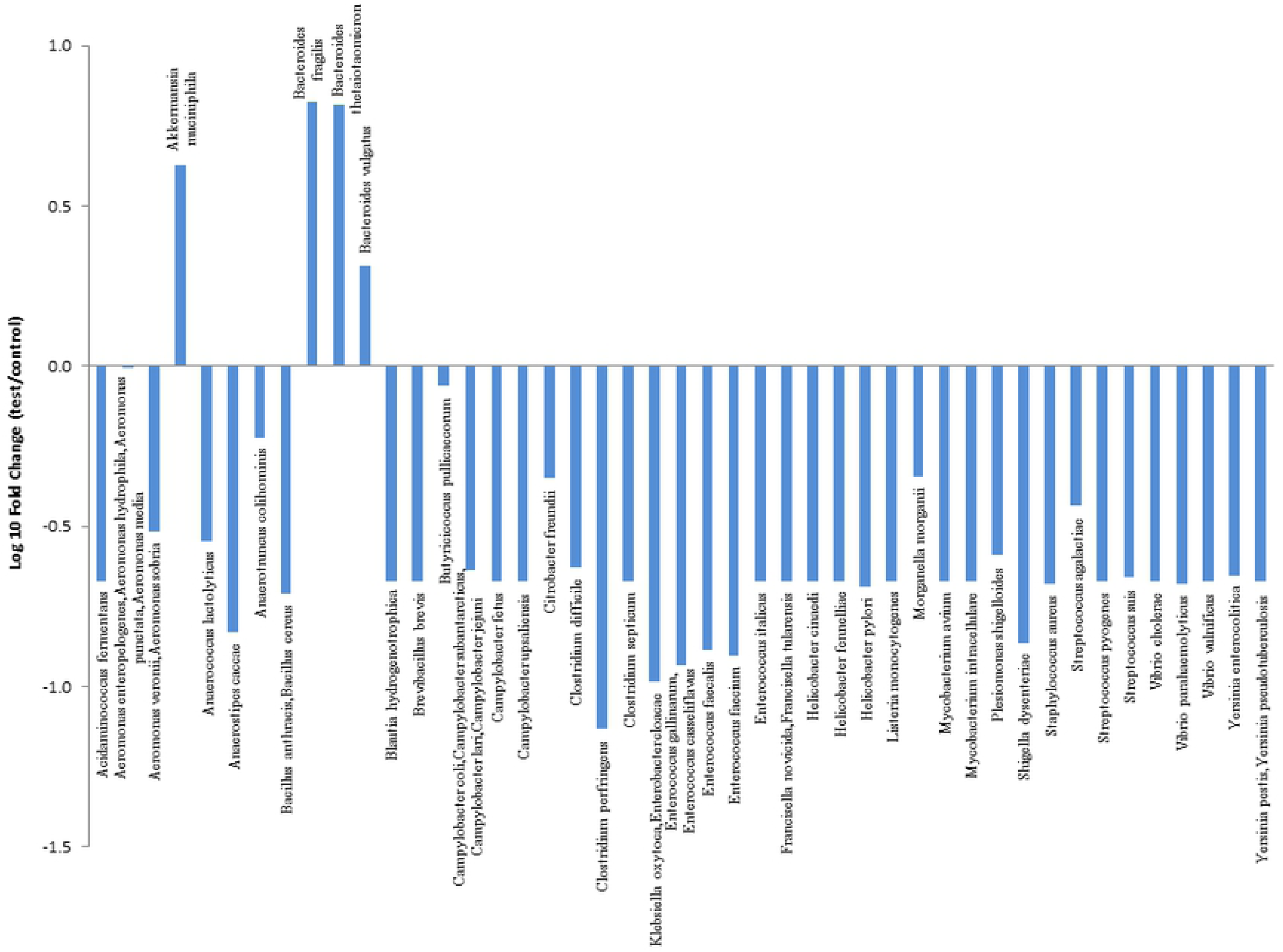
Gut microbial patterns of patients with CRC compared to healthy individuals. The relative abundance of 45 gut microbiota species in patients with CRC (n=24) and healthy individuals (n=44) were investigated using a PCR array amplifying the 16S rRNA genes in stool samples. The relative abundance of each microbial species relative to the abundance in healthy volunteers is shown.

### Changes in the patterns of gut microbiota in patients with diabetes mellitus when compared to healthy individuals

The relative abundance of the gut microbiota has shown to be different in those with metabolic diseases such as DM [7, 18]. As 17/24 (70.83%) patients with CRC also had DM, we proceed to determine if these changes in the gut microbiota observed in those with CRC were associated with CRC or with DM. In order to determine the changes associated with DM, we compared the patterns of gut microbiota in patients with DM (n= 20) with the group of healthy volunteers (n=44). Again. we observed marked differences in the gut microbiota pattern in patients with DM compared to healthy individuals (Fig 2).

**Figure 2:**
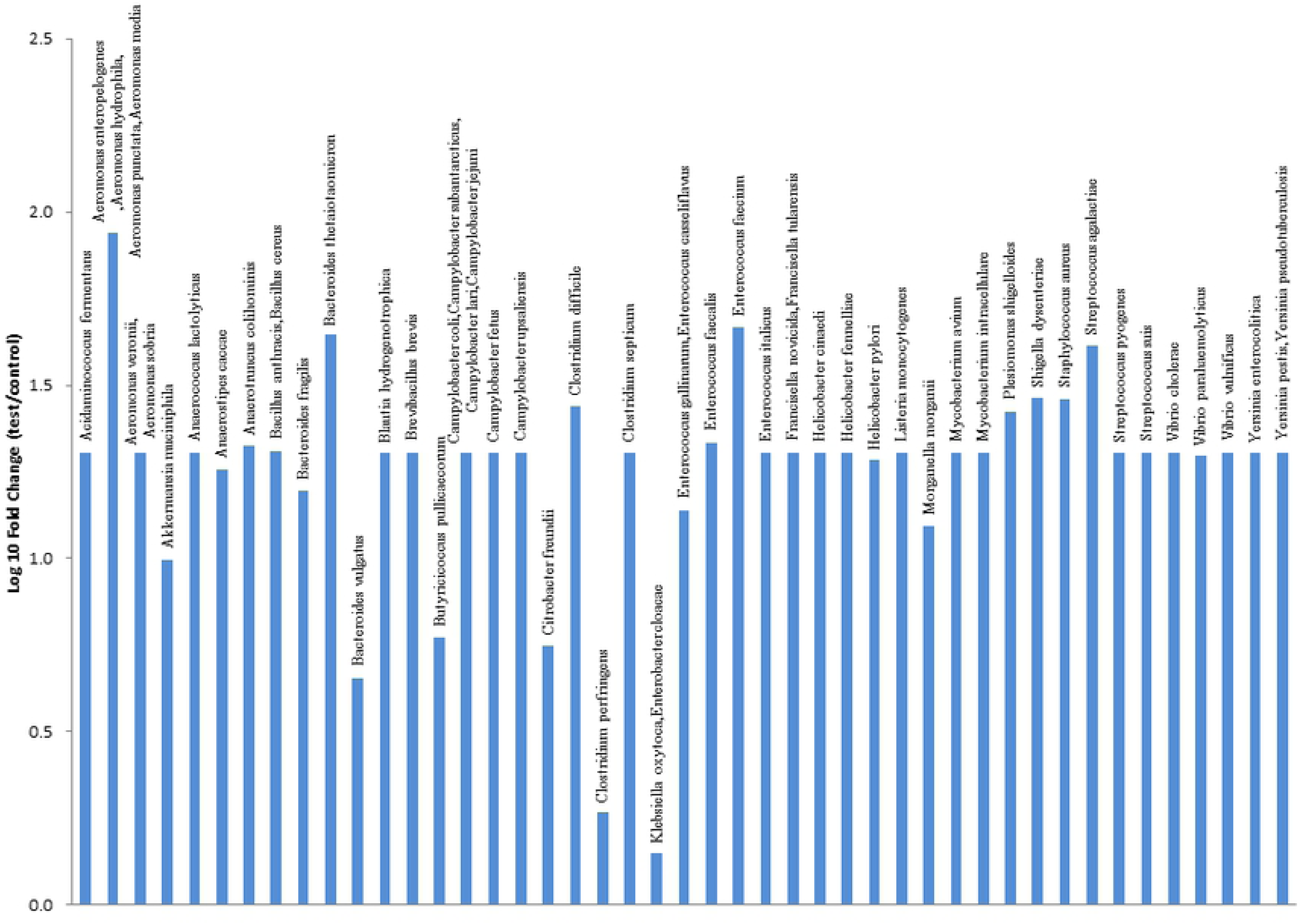
Gut microbial patterns of patients with DM compared to healthy individuals. The relative abundance of 45 gut microbiota species in patients with DM (n=20) and healthy individuals (n=44) were investigated using a PCR array amplifying the 16S rRNA genes in stool samples. The relative abundance of each microbial species relative to the abundance in healthy volunteers is shown.

*Aeromonas spp. (Aeromonas enteropelogenes, Aeromonas hydrophila, Aeromonas punctate, Aeromonas media*) were the predominant microbes, in patients with DM, which were seen at 226.64-fold times higher than healthy individuals. The abundance of *Enterococcus faecium, Bacteroides thetaiotaomicron, Streptococcus agalactiae. Shigella dysenteriae, Enterococcus faecalis, Bacteroides fragilis and Plesiomonas shigelloides* were also several fold higher in the stool samples of DM patients compared to healthy individuals (Table 2) followed by *E. faecium, S. dysenteriae* and *Streptococcus agalactiae.* In contrast, to what we observed in patients with CRC, all the 45 bacterial species investigated several folds more abundant in patients with DM compared to healthy individuals.

### Comparison of the gut microbial patterns of CRC patients compared to those with DM

As we observed marked differences the relative abundance of the gut microbiota between those with CRC and healthy individuals and those with DM and healthy individuals, and since 17/24 (70.83%) of the individuals with CRC had DM, we proceeded to analyze the gut microbial patterns specific to CRC by comparing the microbial patterns of those with CRC (n=24) with those with DM (n=20).

Although all 45 bacterial species were overabundant in patients with DM compared to those with CRC, of whom 70.8% had DM, *Bacteroides fragilis, thetaiotaomicron* and *Akkermansia muciniphila* were least expressed (Figure 3). As, these 3 species of bacteria appear to be associated with CRC, we evaluated the expression of these 3 species related to tumour grade and site of the tumor. There was no difference in the abundance of these 3 species of bacteria with the stage 0 to ii (n=9), iii (n=9) and iv (n=6), when analysed using the Kruskal-Wallis test. There was also no difference in their abundance based on gender or age.

**Figure 3:**
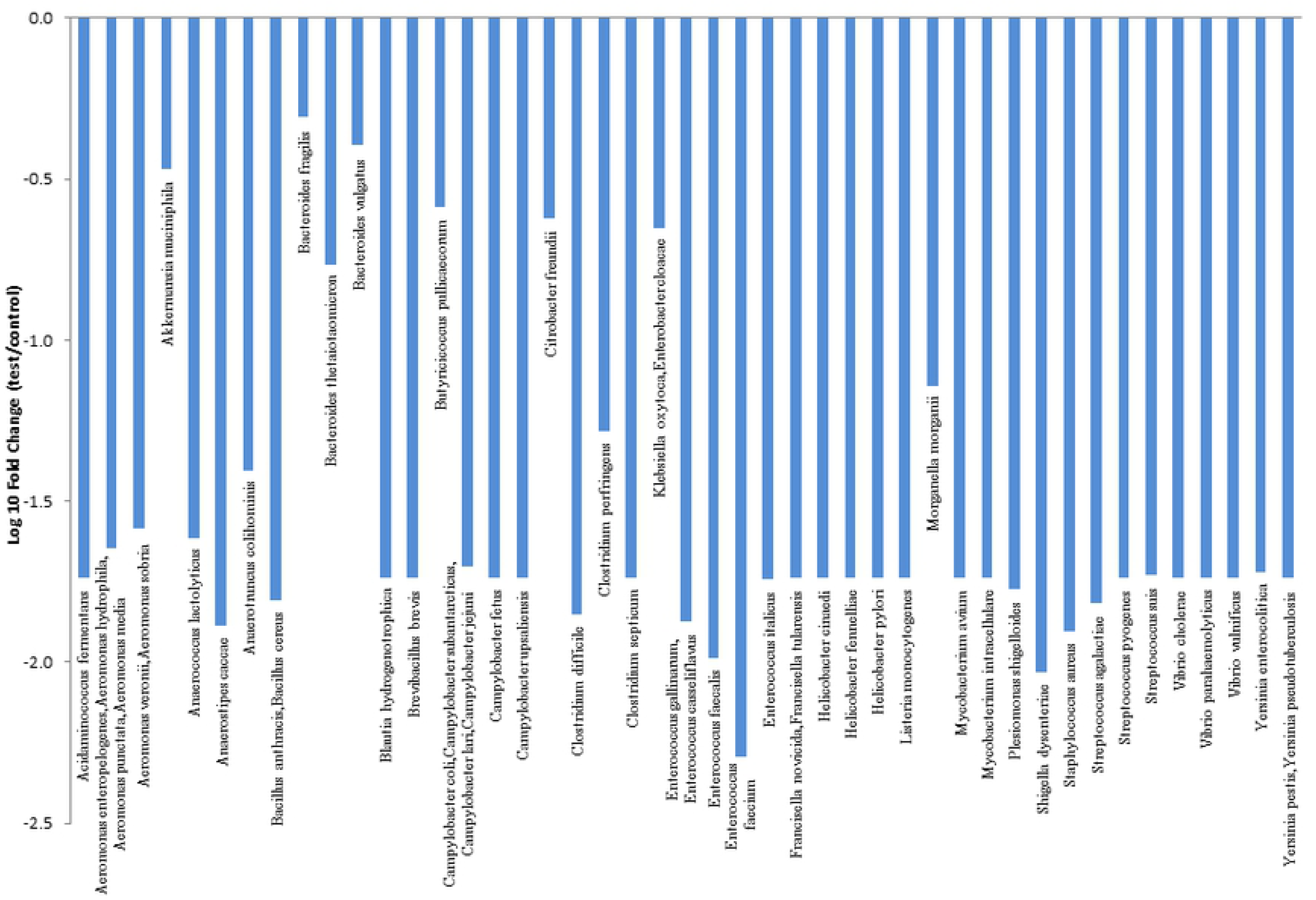
Gut microbial patterns of patients with CRC compared to patients with DM. The relative abundance of 45 gut microbiota species in patients with CRC (n=24) and patients with diabetes mellitus (n=20) were investigated using a PCR array amplifying the 16S rRNA genes in stool samples. The relative abundance of each microbial species relative to the abundance in healthy volunteers is shown.

## Discussion

In this study we investigated the differences in the gut microbial patterns in a South Asian cohort of individuals with CRC and found that the gut microbial patterns were vastly different between those with CRC and healthy individuals. As a large proportion (70.83%) of those with CRC also had diabetes, in order to determine if the changes in the relative abundance of different bacteria were due to the presence of diabetes, we compared the gut microbial patterns in patients with DM with age matched healthy individuals. Again, we found that the gut microbial patterns were indeed markedly different in those with DM compared to healthy individuals. Further comparison of the microbial patients with DM with those with CRC showed that three bacterial species were more likely to be associated with CRC. These are *Bacteroides fragilis, Bacteroides thetaiotaomicron* and *Akkermansia muciniphila.* However, the relative abundance of these three bacterial species did not differ based on the tumour stage, age or the gender of patients.

Many previous studies have shown the association between CRC and *Bacteroides fragilis* [19]. The toxin produced by Enterotoxigenic *Bacteroides fragilis* has shown to change gene transcription in the colons of mice models by inducing epigenetic changes, which subsequently result in development of tumors [20]. The toxin producing strains of *Bacteroides fragilis* has shown to be more prevalent in patients with CRC [10]. Although we did not specifically assess toxin production by *Bacteroides fragilis* in this study, this bacterium was found to be 23.9 times overabundant in those with CRC compared to healthy individuals. Although we found that *Bacteroides vulgatus* was also two-fold over abundant in those with CRC compared to healthy individuals, other studies have shown that *Bacteroides vulgatus* was more abundant in healthy individuals [21]. In addition, we found that *Bacteroides thetaiotaomicron* was 8-fold more over abundant in those with CRC when compared to those with CRC, which has not been reported before.

Ethnic differences have shown to play a significant role in the outcome of CRC, which has been attributed to possible differences in the gut microbiome [22]. For instance, while Bacteroides species, *Fusobacterium nucleatum* and *Enterobacter* species were more abundant in African-American patients with CRC, *Akkermansia muciniphila* and *Bifidobacterium* species were more abundant in Caucasians [22]. As our PCR array did not have *Fusobacterium nucleatum* and *Bifidobacterium* species included, we could not assess their relative abundance. However, *Akkermansia muciniphila* was 5.9-fold over abundant in those with CRC, compared to healthy individuals. Therefore, it would be important to carry out 16S sequencing of the whole gut microbiome to derive better data to find out the microbiota that associate with CRC in the Sri Lankan population.

As 70.8% of the CRC patients also had DM, in order to identify the gut microbiota patterns related to DM, we assessed the patterns between healthy individuals and those with DM. We found that all the 45 types of gut microbiota assessed in this study, were several folds over expressed in those with DM compared to healthy individuals. Specifically, the Aeromonas species, *Enterococcus faecium* and *Shigella dysenteriae* were over 100-fold over abundant in those with DM compared to healthy individuals. Therefore, bacteria of the Phylum Proteobacteria do appear to be 100-fold more abundant in South Asian patients with DM, similar to the observations in Western countries, despite differences in their diets [23]. This gut microbial dysbiosis that occurs due to the overgrowth of bacteria of the Phylum Proteobacteria has shown to associate with low grade endotoxaemia [24]. The presence of low levels of bacterial lipopolysaccharide (LPS) has shown to associate with DM and other metabolic diseases such as non-alcoholic steatohepatitis, which are rapidly increasing in all South Asian countries [25-27]. Therefore, similar patterns of gut microbial dysbiosis that associate with CRC and DM appear be seen in South Asian populations as in other countries, despite differences in the diet and ethnicity.

In summary, we studied the gut microbial patterns in a South Asian cohort of patients with CRC and found that *Bacteroides fragilis, Bacteroides thetaiotaomicron* and *Akkermansia muciniphila* were several folds over abundant in those with CRC when compared to healthy individuals. However, as this study was limited to studying only 45 genera and species of microbiota, it would be important to study the whole gut microbiome by carrying out 16S sequencing to identify other possible microbes that associate with CRC. Early identification of such gut microbial dysbiosis could lead to prevention and treatment strategies in populations by possible nutrition interventions.

## Supporting information captions

Supplementary table 1: The Microbial DNA qPCR Array Intestinal Infection 2 kit (Qiagen, Hilden, Germany)

## References

1. Araghi M, Soerjomataram I, Jenkins M, Brierley J, Morris E, Bray F, et al. Global trends in colorectal cancer mortality: projections to the year 2035. Int J Cancer. 2018. doi: 10.1002/ijc.32055. PubMed PMID: 30536395.

2. Tsoi KKF, Hirai HW, Chan FCH, Griffiths S, Sung JJY. Predicted Increases in Incidence of Colorectal Cancer in Developed and Developing Regions, in Association With Ageing Populations. Clin Gastroenterol Hepatol. 2017;15(6):892–900 e4. doi: 10.1016/j.cgh.2016.09.155. PubMed PMID: 27720911.

3. Cornish AJ, Tomlinson IPM, Houlston RS. Mendelian randomisation: A powerful and inexpensive method for identifying and excluding non-genetic risk factors for colorectal cancer. Mol Aspects Med. 2019. doi: 10.1016/j.mam.2019.01.002. PubMed PMID: 30710596.

4. Banjari I, Kozic S. Dietary intake of vitamin B12 in relation to diet and lifestyle characteristics in a population at high risk for colorectal cancer. Cent Eur J Public Health. 2018;26(4):253–9. doi: 10.21101/cejph.a4585. PubMed PMID: 30660133.

5. Ballotari P, Vicentini M, Manicardi V, Gallo M, Chiatamone Ranieri S, Greci M, et al. Diabetes and risk of cancer incidence: results from a population-based cohort study in northern Italy. BMC Cancer. 2017;17(1):703. doi: 10.1186/s12885-017-3696-4. PubMed PMID: 29070034; PubMed Central PMCID: PMCPMC5657107.

6. Beaugerie L, Svrcek M, Seksik P, Bouvier AM, Simon T, Allez M, et al. Risk of colorectal high-grade dysplasia and cancer in a prospective observational cohort of patients with inflammatory bowel disease. Gastroenterology. 2013;145(1):166–75 e8. doi: 10.1053/j.gastro.2013.03.044. PubMed PMID: 23541909.

7. Kho ZY, Lal SK. The Human Gut Microbiome - A Potential Controller of Wellness and Disease. Front Microbiol. 2018;9:1835. doi: 10.3389/fmicb.2018.01835. PubMed PMID: 30154767; PubMed Central PMCID: PMCPMC6102370.

8. Shimpoh T, Hirata Y, Ihara S, Suzuki N, Kinoshita H, Hayakawa Y, et al. Prevalence of pks-positive Escherichia coli in Japanese patients with or without colorectal cancer. Gut Pathog. 2017;9:35. doi: 10.1186/s13099-017-0185-x. PubMed PMID: 28616082; PubMed Central PMCID: PMCPMC5468999.

9. Bonnet M, Buc E, Sauvanet P, Darcha C, Dubois D, Pereira B, et al. Colonization of the human gut by E. coli and colorectal cancer risk. Clinical cancer research : an official journal of the American Association for Cancer Research. 2014;20(4):859–67. doi: 10.1158/1078-0432.CCR-13-1343. PubMed PMID: 24334760.

10. Boleij A, Hechenbleikner EM, Goodwin AC, Badani R, Stein EM, Lazarev MG, et al. The Bacteroides fragilis toxin gene is prevalent in the colon mucosa of colorectal cancer patients. Clin Infect Dis. 2015;60(2):208–15. doi: 10.1093/cid/ciu787. PubMed PMID: 25305284; PubMed Central PMCID: PMCPMC4351371.

11. Flanagan L, Schmid J, Ebert M, Soucek P, Kunicka T, Liska V, et al. Fusobacterium nucleatum associates with stages of colorectal neoplasia development, colorectal cancer and disease outcome. European journal of clinical microbiology & infectious diseases : official publication of the European Society of Clinical Microbiology. 2014;33(8):1381–90. doi: 10.1007/s10096-014-2081-3. PubMed PMID: 24599709.

12. Allen J, Sears CL. Impact of the gut microbiome on the genome and epigenome of colon epithelial cells: contributions to colorectal cancer development. Genome Med. 2019;11(1):11. doi: 10.1186/s13073-019-0621-2. PubMed PMID: 30803449.

13. Moreno-Indias I, Cardona F, Tinahones FJ, Queipo-Ortuno MI. Impact of the gut microbiota on the development of obesity and type 2 diabetes mellitus. Front Microbiol. 2014;5:190. doi: 10.3389/fmicb.2014.00190. PubMed PMID: 24808896; PubMed Central PMCID: PMCPMC4010744.

14. De Filippis F, Pellegrini N, Vannini L, Jeffery IB, La Storia A, Laghi L, et al. High-level adherence to a Mediterranean diet beneficially impacts the gut microbiota and associated metabolome. Gut. 2016;65(11):1812–21. doi: 10.1136/gutjnl-2015-309957. PubMed PMID: 26416813.

15. Deschasaux M, Bouter KE, Prodan A, Levin E, Groen AK, Herrema H, et al. Depicting the composition of gut microbiota in a population with varied ethnic origins but shared geography. Nature medicine. 2018;24(10):1526–31. doi: 10.1038/s41591-018-0160-1. PubMed PMID: 30150717.

16. Louis P, Hold GL, Flint HJ. The gut microbiota, bacterial metabolites and colorectal cancer. Nature reviews. 2014;12(10):661–72. doi: 10.1038/nrmicro3344. PubMed PMID: 25198138.

17. Compton C, Fenoglio-Preiser CM, Pettigrew N, Fielding LP. American Joint Committee on Cancer Prognostic Factors Consensus Conference: Colorectal Working Group. Cancer. 2000;88(7):1739–57. PubMed PMID: 10738234.

18. Dugas LR, Lie L, Plange-Rhule J, Bedu-Addo K, Bovet P, Lambert EV, et al. Gut microbiota, short chain fatty acids, and obesity across the epidemiologic transition: the METS-Microbiome study protocol. BMC public health. 2018;18(1):978. doi: 10.1186/s12889-018-5879-6. PubMed PMID: 30081857; PubMed Central PMCID: PMCPMC6090745.

19. Drewes JL, White JR, Dejea CM, Fathi P, Iyadorai T, Vadivelu J, et al. High-resolution bacterial 16S rRNA gene profile meta-analysis and biofilm status reveal common colorectal cancer consortia. NPJ Biofilms Microbiomes. 2017;3:34. doi: 10.1038/s41522-017-0040-3. PubMed PMID: 29214046; PubMed Central PMCID: PMCPMC5707393.

20. Allen J, Hao S, Sears CL, Timp W. Epigenetic changes induced by Bacteroides fragilis toxin (BFT). Infection and immunity. 2019. doi: 10.1128/IAI.00447-18. PubMed PMID: 30885929.

21. Wang T, Cai G, Qiu Y, Fei N, Zhang M, Pang X, et al. Structural segregation of gut microbiota between colorectal cancer patients and healthy volunteers. ISME J. 2012;6(2):320–9. doi: 10.1038/ismej.2011.109. PubMed PMID: 21850056; PubMed Central PMCID: PMCPMC3260502.

22. Farhana L, Antaki F, Murshed F, Mahmud H, Judd SL, Nangia-Makker P, et al. Gut microbiome profiling and colorectal cancer in African Americans and Caucasian Americans. World J Gastrointest Pathophysiol. 2018;9(2):47–58. doi: 10.4291/wjgp.v9.i2.47. PubMed PMID: 30283710; PubMed Central PMCID: PMCPMC6163128.

23. Shin NR, Whon TW, Bae JW. Proteobacteria: microbial signature of dysbiosis in gut microbiota. Trends Biotechnol. 2015;33(9):496–503. doi: 10.1016/j.tibtech.2015.06.011. PubMed PMID: 26210164.

24. Hawkesworth S, Moore SE, Fulford AJ, Barclay GR, Darboe AA, Mark H, et al. Evidence for metabolic endotoxemia in obese and diabetic Gambian women. Nutr Diabetes. 2013;3:e83. doi: 10.1038/nutd.2013.24. PubMed PMID: 23978817; PubMed Central PMCID: PMCPMC3759130.

25. Cani PD, Osto M, Geurts L, Everard A. Involvement of gut microbiota in the development of low-grade inflammation and type 2 diabetes associated with obesity. Gut Microbes. 2012;3(4):279–88. doi: 10.4161/gmic.19625. PubMed PMID: 22572877; PubMed Central PMCID: PMCPMC3463487.

26. Wigg AJ, Roberts-Thomson IC, Dymock RB, McCarthy PJ, Grose RH, Cummins AG. The role of small intestinal bacterial overgrowth, intestinal permeability, endotoxaemia, and tumour necrosis factor alpha in the pathogenesis of non-alcoholic steatohepatitis. Gut. 2001;48(2):206–11. PubMed PMID: 11156641; PubMed Central PMCID: PMCPMC1728215.

27. Pussinen PJ, Havulinna AS, Lehto M, Sundvall J, Salomaa V. Endotoxemia Is Associated With an Increased Risk of Incident Diabetes. Diabetes care. 2011;34(2):392–7. doi: 10.2337/dc10-1676.

